# Contrastive Learning for Omics-guided Whole-slide Visual Embedding Representation

**DOI:** 10.1101/2025.01.12.632280

**Authors:** Suwan Yu, Yooeun Kim, Hoeyoung Kim, Sangseon Lee, Kwangsoo Kim

## Abstract

While computational pathology has transformed cancer diagnosis and prognosis prediction, existing computational methods remain limited in their ability to decipher the complex molecular characteristics within tumors. We present CLOVER (Contrastive Learning for Omics-guided whole-slide Visual Embedding Representation), a novel deep learning framework that leverages self-supervised contrastive learning to integrate multi-omics data (genomics, epigenomics, and transcriptomics) with slide representations, connecting the morphological and molecular features of tumors. Using the breast cancer cohorts comprising diagnostic slides and multi-omics paired data from 610 breast cancer patients, we validated CLOVER’s excellence by demonstrating its ability to generate effective slide-level representations that consider molecular states of cancer. CLOVER outperforms existing methods in few-shot learning scenarios, particularly in cancer subtype classification and clinical biomarker prediction tasks (ER, PR, and HER2 status). Through comprehensive interpretability analysis, we identified tumor microenvironment components within slides and revealed molecular features associated with breast cancer. Our results demonstrate that CLOVER enables detailed molecular characterization from single slide analysis, suggesting its potential for effective utilization in future cancer diagnosis and prognosis prediction studies.

## 1 Introduction

### 1.1 Background of the study

Whole Slide Image (WSI) are fundamental to pathologic diagnosis, providing detailed visual insights into tissue morphology, location, and cellular distribution at the microscopic level [1]. These high-resolution digital scans allow pathologists to assess critical morphological features necessary for determining malignancy, classifying subtypes, and predicting patient responses to therapy [2,3]. However, the immense size and complexity of WSI—often containing billions of pixels per image—make manual analysis time-consuming, labor-intensive, and subject to inter-observer variability, highlighting the pressing need for automated and reproducible computational methods [4].

Digital pathology has transformed cancer diagnosis and prognosis prediction through computational methods that can extract complex patterns directly from image data, enabling sophisticated tasks such as survival prediction, prognosis assessment, and treatment response estimation [5]. Nevertheless, the development and deployment of these computational models face significant challenges due to the labor-intensive nature of data collection and annotation [6], particularly when developing specialized models for the vast array of diagnostic categories and rare diseases encountered in clinical practice.

### 1.2 Current challenges in computational pathology

Self-supervised learning (SSL) has emerged as a promising solution to address the annotation scarcity problem by allowing models to derive meaningful representations from unlabeled data [7]. While these approaches have shown success in capturing morphological patterns, they are inherently limited by their reliance on visual features alone, potentially missing crucial molecular characteristics that influence disease progression. Recent advances in multimodal learning have attempted to bridge this gap by integrating additional data modalities with images [8–12]. However, current methods face several limitations: (1) Vision-language models, while improving interpretability, cannot capture underlying molecular mechanisms; (2) Single-omics integration approaches might be insufficient to represent the complex molecular interactions in tumor biology; and (3) Existing methods face challenges in effectively combining heterogeneous data sources while maintaining biological interpretability.

### 1.3 Purpose of the research

To address these limitations, we developed CLOVER (Contrastive Learning for Omics-guided whole-slide Visual Embedding Representation), an innovative deep learning architecture that leverages SSL to unify multi-omics data into the slide representations. CLOVER integrates diverse molecular data layers, including genomics (copy number variation), epigenomics (DNA methylation), and transcriptomics (mRNA abundances), to guide the learning of image representations. This multi-omics integration allows CLOVER to encapsulate various layers of biological data, addressing the complex molecular mechanisms underlying cancer that single-omics approaches cannot fully capture. Our experiments demonstrate that CLOVER achieves superior performance in few-shot classification scenarios for tasks such as cancer subtype classification and clinical marker status prediction in breast cancer. Through multi-omics guided representation learning, CLOVER enables detailed molecular characterization from single slide analysis without any label, establishing a more comprehensive understanding of tumor biology at both morphological and molecular levels.

## 2 Related Work

### 2.1 Computational pathology

Computational pathology has become integral to cancer diagnosis and prognosis, enabled by large-scale public databases [13–15]. In this field, analysis tasks are typically categorized by their granularity. Patch-level analysis examines small tissue sections (e.g., 256×256 pixels) for specific tasks including tumor region localization [16,17], tissue type classification [18], and nuclei detection and classification [19]. In contrast, slide-level analysis addresses broader diagnostic and prognostic tasks such as cancer subtype classification [20] and survival prediction [21].

The extremely high dimensionality of WSI poses significant challenges for traditional deep learning models such as Convolutional Neural Networks (CNN) or Vision Transformers (ViTs) [22]. Multiple Instance Learning (MIL) has emerged as the preferred framework for slide-level analysis, treating each WSI as a bag containing multiple instances (patches) [23–25]. In MIL-based approaches, supervision is only available at the bag level, and the model must learn to aggregate information by attention mechanisms that assign importance weights to different patches based on their relevance. Recent MIL architectures, such as Attention-based MIL (ABMIL) [26] and TransMIL [27], have demonstrated particular success in capturing long-range dependencies between patches while maintaining computational efficiency.

### 2.2 Image-based self-supervised learning in pathology

Self-supervised learning (SSL) in pathology have evolved significantly with the emergence of foundation models, typically defined as those trained on more than 100,000 WSIs [28–31]. SSL is a machine learning approach that creates training labels from unlabeled data itself, allowing models to learn useful features without manual annotations. These approaches operate at both patch and slide levels, employing diverse learning strategies. At the patch level, models adopt advanced self-supervised learning strategies to capture meaningful representations. UNI [28] and Virchow [30] leverage self-distillation with DINOv2 [32] framework, which focuses on semantic alignment through teacher-student learning. Also, BEPH [33] employ masked image modeling using BEiT [34] framework, emphasizing the reconstruction of masked regions to capture fine-grained tissue structures and morphology.

Slide-level SSL approaches address the additional challenge of information aggregation across patches. Models such as HIPT [35] introduce hierarchical learning architectures that build representations at multiple scales, while Prov-GigaPath [36] leverage sophisticated attention mechanisms for whole-slide feature integration. These foundation models, trained on diverse tissue types, have demonstrated strong transfer learning capabilities across various pathological tasks, including cancer detection, subtype classification, and prognosis prediction. However, they remain limited by their focus on image features alone.

### 2.3 Multimodal learning in pathology

Given that pathology practice heavily relies on natural language for patient reports, academic publications, and medical education, Vision-Language Models (VLMs) in pathology have emerged in two main directions: report-based learning and caption-based learning, building upon successful architectures like CLIP [37] and CoCa [38]. Models like PRISM [39] learn from detailed pathology reports, while approaches such as CONCH [40] utilize more concise image descriptions to enhance model interpretability. These VLM approaches enable diverse applications, from automated report generation and image-text retrieval to zero-shot disease classification and diagnostic support.

More recent work has explored molecular data integration through Vision-Omics models. TANGLE pioneered this direction by aligning slide image features with transcriptomic profiles through contrastive learning, demonstrating improved cancer subtype classification [41]. However, single-omics approaches might miss the complex interactions between different molecular layers that characterize tumor biology. The integration of multiple omics modalities (genomics, epigenomics, transcriptomics) with WSI analysis represents a crucial frontier in computational pathology [21]. Success in this area requires novel architectural designs that can effectively combine these diverse data types while preserving their biological relationships and yielding interpretable predictions.

## 3 Methodology

Here, we present CLOVER (Contrastive Learning for Omics-guided Visual Embedding Representation), a framework designed to learn shared representations between whole slide images and multi-omics data. CLOVER consists of three main components: (1) a slide encoder that processes histological images into slide-level embeddings, (2) a multi-omics encoder that integrates multiple molecular data types into a unified representation, and (3) a multimodal contrastive loss module that learns to bridge these distinct data modalities (Figure 1). Through self-supervised pretraining, the slide encoder learns to capture molecular characteristics in its visual representations, which are then evaluated through few-shot classification tasks on molecular subtypes and characteristics.

**Fig. 1.**
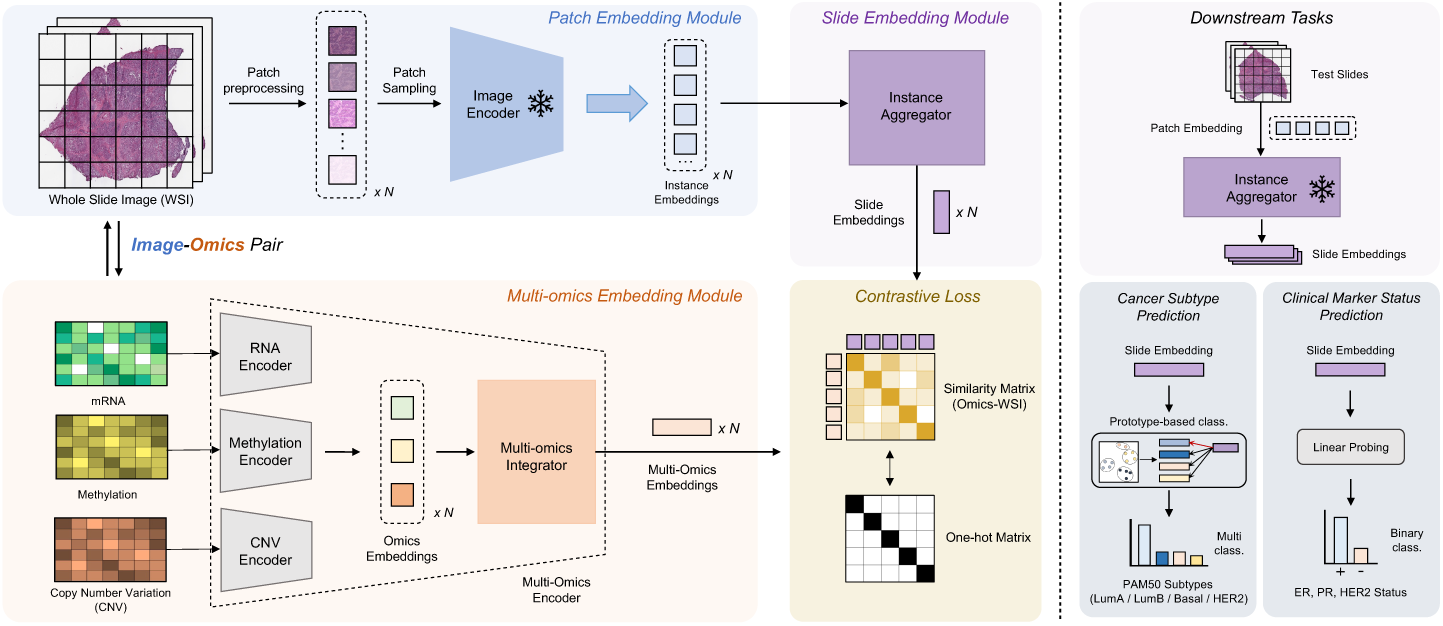
CLOVER overview for pre-training and downstream tasks. The framework integrates histopathological whole slide images (WSI) with multi-omics data (mRNA, Methylation, and CNV) through three main modules: Patch Embedding, Slide Embedding, and Multi-omics Embedding. The model employs contrastive learning between image-omics pairs and can be applied to downstream tasks including cancer subtype prediction and clinical marker status prediction, with frozen components indicating pre-trained elements.

### 3.1 Whole slide image preprocessing

To process whole-slide images (WSIs) for downstream analysis, we implemented a custom patch slicing method followed by background filtering process. The algorithm takes each WSI file as input and extracts image tiles at a predefined magnification level. First, the base magnification of the slide is determined from the slide metadata. The target magnification level is then calculated by dividing the base magnification by the desired magnification (20× in this study). The patch gets sliced with the size of 256 pixels * 256 pixels and no overlap between patches. The algorithm iterates over all patches and calculates an edge score for each tile using an edge detection filter. Patches with edge scores below a defined threshold (15 in this study) are filtered out. This ensures that only tiles with significant structural features are retained, optimizing storage and downstream computational efficiency. We applied an additional filtering process using Hover-Net [42] extracted statistics. At the patch level, patches where the majority of cell annotations were classified as ‘no-neo’ (no-neoplastic) or lacked labels were excluded. Patches with fewer than five cells were also removed to ensure sufficient information. Among the remaining patches, those with a major cell type proportion, determined by majority voting, below 0.3 were further excluded.

### 3.2 Model architecture

#### Slide encoder

We adopt a hierarchical encoding strategy following the Multiple Instance Learning (MIL) paradigm. The slide encoder comprises two key components:

##### Patch Image Encoder

We utilize UNI[28], a state-of-the-art foundation model pretrained on over 100 million patches from 100,000+ diagnostic H&E-stained WSIs. For the *i*-th slide, we denote the resulting patch embeddings as 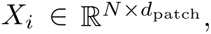 where *N* is the number of patches (*N* = 2, 048) and *d*_patch_ is the embedding dimension (*d*_patch_ = 1, 024). These embeddings are then processed through a projection layer before being fed into the instance aggregator.

##### Instance Aggregator

To aggregate patch-level information into a slide-level representation, we employ an Attention-Based Multiple Instance Learning (ABMIL) model [26]. The aggregator function 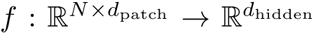 maps patch embeddings to a slide embedding 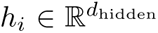 through learned attention weights, where *d*_hidden_ = 786.

#### Multi-omics encoder

The multi-omics encoder processes three distinct molecular data types: RNA sequencing, DNA methylation, and copy number variation. Each modality is encoded through a dedicated two-layer Self-Normalizing Network (SNN), generating 256-dimensional embeddings. These modality-specific embeddings are concatenated and projected through a multi-omics aggregator to produce a final 786-dimensional representation 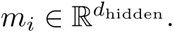

#### Multi-modal contrastive loss

We align the embedding spaces of slide and multi-omics representations using a symmetric cross-modal contrastive learning objective [37]. For a batch of *M* pairs 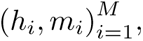 where *h_i_* and *m_i_* are the slide and multi-omics embeddings respectively, the loss function is defined as:

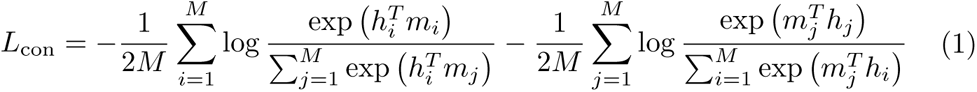

This objective maximizes the similarity between matched pairs while minimizing similarity between unmatched pairs.

### 3.3 Model interpretability

#### Slide-level analysis

Our framework’s attention mechanism identifies regions of high importance in histological images during the embedding generation process. This allows us to extract and analyze patches with high attention scores, revealing diagnostically significant regions within the slides. Additionally, using the Hover-Net [42] pre-trained on PanNuke [43] dataset, we analyze cell type distributions in high-attention patches, categorizing cells into neoplastic, inflammatory, connective, dead.

#### Omics-level analysis

To understand the molecular features driving our model’s embedding alignment, we apply integrated gradients methodology [44] to analyze feature importance in the multi-omics data. This enables attribution of learned representations to specific molecular features across RNA-seq, DNA methylation, and CNV data, providing insights into the molecular basis of morphological patterns. By examining these highly weighted molecular features and their known biological functions, we can understand what molecular characteristics our model learns to associate with histological patterns during self-supervised training.

## 4 Experiments and Results

### 4.1 Data collection and experimental design

We evaluated our methodology using the The Cancer Genome Atlas Breast Invasive Carcinoma (TCGA-BRCA) cohorts [14], which provides multimodal data from breast cancer patients, including paired whole slide images and multi-omics profiles (copy number variation, DNA methylation, and RNA-seq data) from tumor tissues. The dataset comprises 610 patients, with 658 slides and corresponding multi-omics profiles. We used diagnostic slides from formalin-fixed paraffin-embedded (FFPE) tissues, which represent the standard for clinical diagnosis and prognostic assessment.

To maintain the integrity of multimodal relationships, we adopted a patient-level approach, preventing improper sampling practices that might associate multiple slides with a single omics profile [41]. We curated metadata from the cBioPortal [45] and GDC portals [46], including histologic subtype, molecular subtype (based on PAM50 classification), and estrogen receptor (ER), progesterone receptor (PR), human epidermal growth factor receptor 2 (HER2) status based on immunohistochemistry (IHC).

The histologic subtypes comprised Invasive Ductal Carcinoma (IDC) and Invasive Lobular Carcinoma (ILC), represented by 493 and 117 slides, respectively. The molecular subtypes showed the following distribution: Luminal A (312), Luminal B (116), Basal-like (112), and HER2-enriched (48) slides. For clinical marker status, we extracted slide-level HER2, ER, and PR information from IHC data curated in the GDC portal. After excluding equivocal, indeterminate, and unavailable cases, we retained only Positive and Negative classifications, resulting in 74/308 slides for HER2 (positive/negative), 433/143 for ER (positive/negative), and 433/143 for PR (positive/negative).

The preprocessed omics data was obtained from previous article, which established pan-cancer normalization and feature selection protocols [47,48]. We used three types of omics data: RNA (RNA-seq), MET (DNA Methylation Array), and CNV (Copy Number Variation). As described in their work, RNA-seq data were normalized using a log2(TPM + 1) transformation, and genes with a standard deviation greater than one were retained, resulting in 6,016 features. DNA methylation preprocessing excluded probes with beta-values below 0.3 in more than 90% of samples, leaving 6,617 informative features for analysis. Copy number variation (CNV) data were filtered by removing segments with less than 5% zeros, a mean absolute value of 0.20 or lower, or a coefficient of variation of 0.20 or lower. This filtering process resulted in the retention of 7,460 informative CNV features.

For evaluation, we employed two distinct evaluation strategies. For representation evaluation, we utilized the entire dataset for pretraining to assess how well the learned slide embeddings capture molecular characteristics, and measured clustering quality between different cancer subtypes. To decide the applicability of the pretraining framework to several cancer classification tasks, we used few-shot classification, employing 5-fold cross-validation to assess model performance across different data subsets. The folds were carefully stratified to ensure even class distribution for downstream tasks, maintaining consistent proportions of histologic subtypes, molecular subtypes, and receptor status across all splits. This stratification was particularly important given the imbalanced nature of our dataset, especially in molecular subtypes and receptor status classes.

### 4.2 Breast cancer slides representation

Accurate representation of histopathological slides is essential for capturing both morphological and molecular characteristics of breast cancer, which are essential for diagnosis, prognosis, and treatment planning. In this study, we evaluated CLOVER’s representation capacity across three critical breast cancer classification criteria: histologic subtypes, molecular subtypes, and triple-negative breast cancer (TNBC) subtype.

Histologic subtype classification, particularly distinguishing between Invasive Ductal Carcinoma (IDC) and Invasive Lobular Carcinoma (ILC), plays a crucial role in determining surgical strategies and therapeutic approaches [49]. IDC typically forms well-defined masses with clear borders, while ILC exhibits diffuse growth patterns that necessitate more extensive surgical margins and demonstrate distinct responses to chemotherapy [50–53]. Accurate differentiation between these subtypes is essential as it directly impacts clinical decision-making. The PAM50 molecular subtype classification system categorizes breast cancer into four subtypes—Luminal A, Luminal B, HER2-enriched, and Basal-like—each with unique prognostic implications and therapeutic responses [54]. For instance, Luminal subtypes respond well to hormone therapy [55], while HER2-enriched tumors benefit from targeted HER2 therapies [56]. Additionally, Basal-like tumors, often overlapping with TNBC, exhibit aggressive clinical behavior and demand tailored therapeutic approaches [57]. TNBC, characterized by the absence of estrogen receptor (ER), progesterone receptor (PR), and HER2 expression, constitutes 15–20% of breast cancers and is associated with poor prognosis [58]. The lack of conventional hormone-directed targets necessitates alternative treatment strategies, including chemotherapy, immune checkpoint inhibitors, and emerging therapies such as PARP inhibitors and antibody-drug conjugates [59]. To evaluate CLOVER’s representation capacity, we trained the model using TCGA-BRCA whole slide images and multi-omics data. We compared CLOVER against two baselines: UNI+Avg, which combines the state-of-the-art UNI model with average pooling for slide-level representations, and TANGLE, which incorporates only RNA-seq data for representation guidance. We assessed representation quality through Uniform Manifold Approximation and Projection (UMAP) visualization and silhouette scores, which quantify the degree of cluster separation between different cancer subtypes.

As shown in UMAP visualizations (Figure 2A), CLOVER consistently produced more distinct and well-separated clusters across all classification criteria compared to baseline methods. For histologic subtype, CLOVER achieved a silhouette score of 0.112, significantly outperforming TANGLE (0.052) and UNI+Avg (0.035). Similarly, for PAM50 molecular subtypes, CLOVER demonstrated superior clustering with a silhouette score of 0.089, compared to TANGLE’s 0.037 and UNI+Avg’s 0.009. In distinguishing TNBC from non-TNBC cases, CLOVER achieved a silhouette score of 0.118, markedly higher than TANGLE (0.052) and UNI+Avg (0.025) (Figure 2B).

**Fig. 2.**
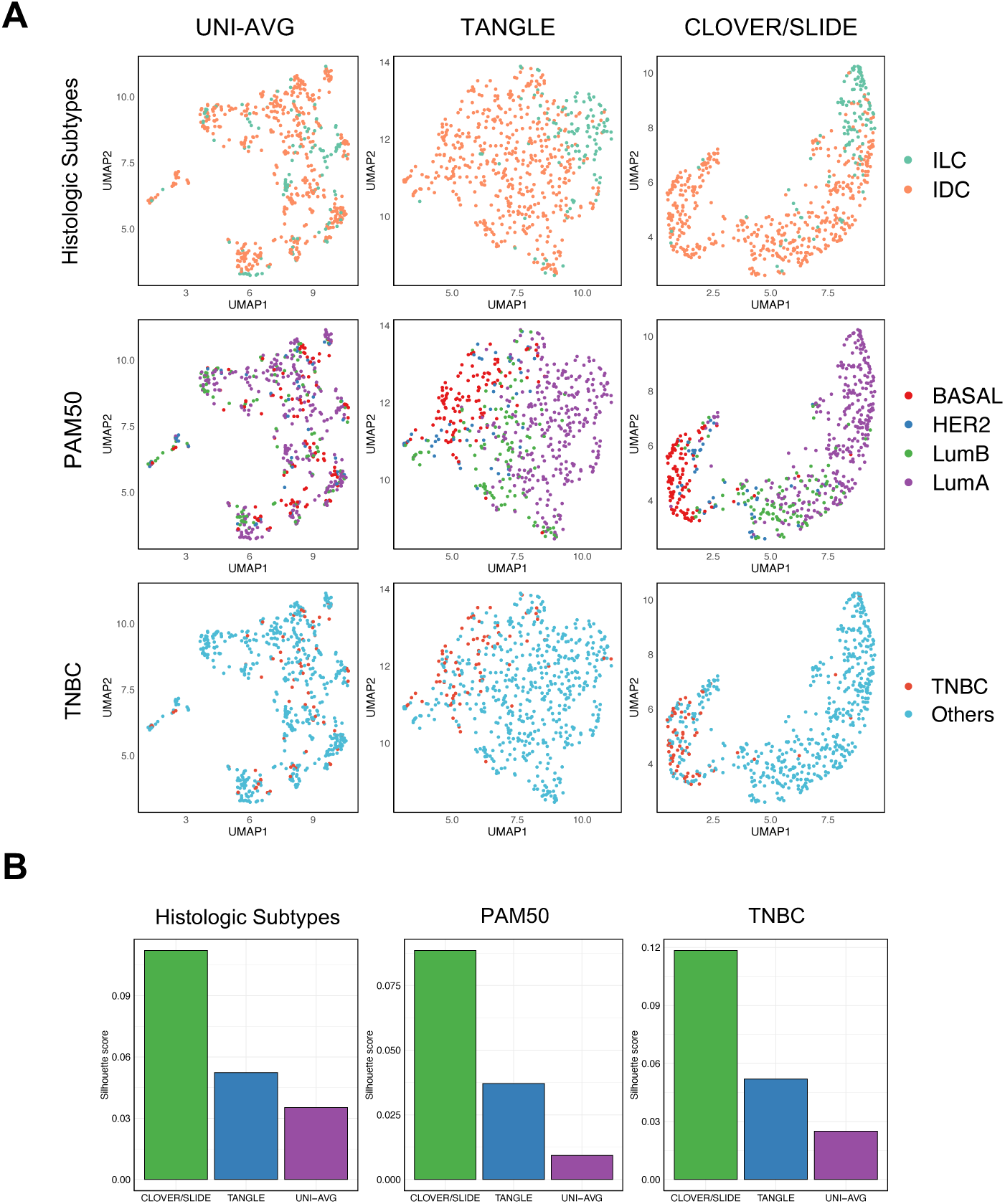
Slide-level representation analysis across breast cancer classification criteria. (A) UMAP visualization comparing embedding spaces generated by UNI+AVG (left), TANGLE (middle), and CLOVER (right) across three classification tasks: histologic subtypes (IDC: blue, ILC: red), PAM50 molecular subtypes (Luminal A: blue, Luminal B: green, Basal-like: red, HER2-enriched: purple), and TNBC subtype (TNBC: red, Non-TNBC: blue). (B) Quantitative comparison using silhouette scores for each classification criterion, demonstrating relative performance of CLOVER (green), TANGLE (navy), and UNI+AVG (purple).

The evaluation of multi-omics integration further highlighted CLOVER’s effectiveness in synthesizing complementary molecular information. Analysis of individual omics modalities revealed varying levels of discriminative power for specific tasks: RNA-seq showed the highest performance for PAM50 classification (silhouette score 0.058), followed by copy number variation (CNV, 0.039) and DNA methylation (0.033). Notably, the integrated multi-omics framework of CLOVER significantly enhanced representation quality, achieving silhouette scores of 0.083 for histologic subtypes, 0.170 for PAM50 classification, and 0.152 for TNBC subtype (Supplementary Figure 1). These results underscore CLOVER’s capacity to integrate diverse molecular modalities, capturing both morphological and molecular tumor characteristics in a biologically meaningful manner.

### 4.3 Few-shot classification performance

We conducted comprehensive few-shot learning experiments to evaluate CLOVER’s ability to learn from limited labeled data across three critical breast cancer classification tasks: histologic subtype classification, PAM50 molecular subtype classification, and clinical marker status prediction. To assess model performance, we used 5-fold cross-validation with 50 random trials per k-shot scenario (k=1,5,10). For each trial, four folds were used for pretraining and one-fold for testing. For binary classification tasks (histologic subtype and clinical markers), we performed linear probing by fixing the pretrained slide encoder and training a logistic regression classifier on the extracted embeddings. For multi-class PAM50 classification, we performed prototype-based classification using learned embeddings, where test samples are classified based on their distance to class prototypes constructed from the limited k-shot examples.

We compared CLOVER against several baseline approaches. For supervised Multiple Instance Learning (MIL) baselines, we implemented UNI+TransMIL and UNI+ABMIL, which combine the UNI patch encoder with TransMIL and AB-MIL aggregation methods respectively. We also evaluated two linear probing baselines: TANGLE, which uses RNA-seq guided embeddings, and UNI+Avg, which applies average pooling to UNI patch embeddings. Performance was assessed using AUROC (Area Under ROC Curve) as the primary metric, supplemented by weighted F1-Score and AUPRC (Area Under the Precision Recall Curve) metrics to account for class imbalance.

#### Histologic subtype classification

In histologic subtype classification, CLOVER demonstrated consistently superior performance across all few-shot scenarios (Table 1). For extreme few-shot conditions (k=1), CLOVER achieved an AU-ROC of 80.7 ± 16.1%, significantly outperforming both TANGLE (77.0 ± 13.7%) and conventional MIL approaches (UNI+ABMIL: 67.6 ± 11.1%, UNI+TransMIL: 65.9 ± 10.7%). This performance advantage extended to scenarios with more training samples, where at k=5, CLOVER achieved an AUROC of 88.1 ± 5.6% compared to TANGLE’s 86.8 ± 5.5%. With k=10 samples per class, CLOVER maintained its superior performance with an AUROC of 89.9 ± 4.8%, demonstrating a 6.9% improvement over the UNI+ABMIL baseline.

**Table 1.** Few-shot Histologic Subtype Classification in Human Breast Cancer. Performance comparison of different models and aggregation methods for classifying breast cancer histologic subtype classification across different few-shot settings (K=1,5,10), evaluated using F1-score, AUROC, and AUPRC metrics.

| Model /<br>Aggregator | K = 1 |  |  | K = 5 |  |  | K = 10 |  |  |
| --- | --- | --- | --- | --- | --- | --- | --- | --- | --- |
|  | F1-score | AUROC | AUPRC | F1-score | AUROC | AUPRC | F1-score | AUROC | AUPRC |
| UNI + | 60.3 ± | 65.9 ± | 36.1 ± | 71.2 ± | 78.9 ± | 53.3 ± | 77.9 ± | 84.6 ± | 63.1 ± |
| TransMIL | 12.9 | 10.7 | 13.6 | 9.2 | 7.6 | 14.1 | 6.2 | 6.0 | 11.6 |
| UNI + | 61.3 ± | 67.6 ± | 38.2 ± | 74.9 ± | 81.2 ± | 57.3 ± | 80.4 ± | 86.7 ± | 67.3 ± |
| ABMIL | 16.3 | 11.1 | 14.3 | 8.4 | 7.4 | 14.2 | 5.7 | 5.7 | 10.9 |
| UNI + | 60.9 ± | 66.4 ± | 36.5 ± | 73.4 ± | 77.9 ± | 51.7 ± | 78.1 ± | 83.0 ± | 60.1 ± |
| Avg | 18.1 | 10.7 | 13.9 | 8.8 | 8.0 | 14.4 | 6.1 | 6.2 | 11.4 |
| TANGLE | 62.9 ± | 77.0 ± | 51.0 ± | 79.5 ± | 86.8 ± | 66.7 ± | 83.8 ± | 89.7 ± | 72.6 ± |
|  | 19.4 | 13.7 | 18.3 | 7.5 | 5.5 | 11.4 | 5.0 | 4.5 | 9.2 |
| CLOVER | **69.7 ±** | **80.7 ±** | **61.6 ±** | **82.3 ±** | **88.1 ±** | **72.4 ±** | **85.3 ±** | **89.9 ±** | **75.8 ±** |
| (Ours) | **15.2** | **16.1** | **21.7** | **7.5** | **5.6** | **12.3** | **5.2** | **4.8** | **10.0** |

The F1-scores and AUPRC metrics further validated CLOVER’s effectiveness, particularly in handling class imbalance between IDC and ILC cases. At k=10, CLOVER achieved an F1-score of 85.3 ± 5.2% and AUPRC of 75.8 ± 10.0%, significantly outperforming all baseline methods. This comprehensive performance improvement suggests that CLOVER’s multi-omics guided representation learning effectively captures subtle morphological differences between histologic sub-types.

#### Molecular subtype classification

For PAM50 molecular subtype classification, CLOVER showed consistent improvements in discriminative power across all few-shot scenarios (Table 2). Starting from k=1, CLOVER achieved a macro-AUROC of 76.7 ± 6.9%, substantially outperforming TANGLE (73.8 ± 6.3%) and UNI+ABMIL (60.1 ± 5.6%). This performance advantage was maintained as training samples increased, with k=5 showing macro-AUROC of 85.8 ± 2.9% and k=10 reaching 87.2 ± 2.2%, both with notably lower variance compared to baseline methods.

**Table 2.**
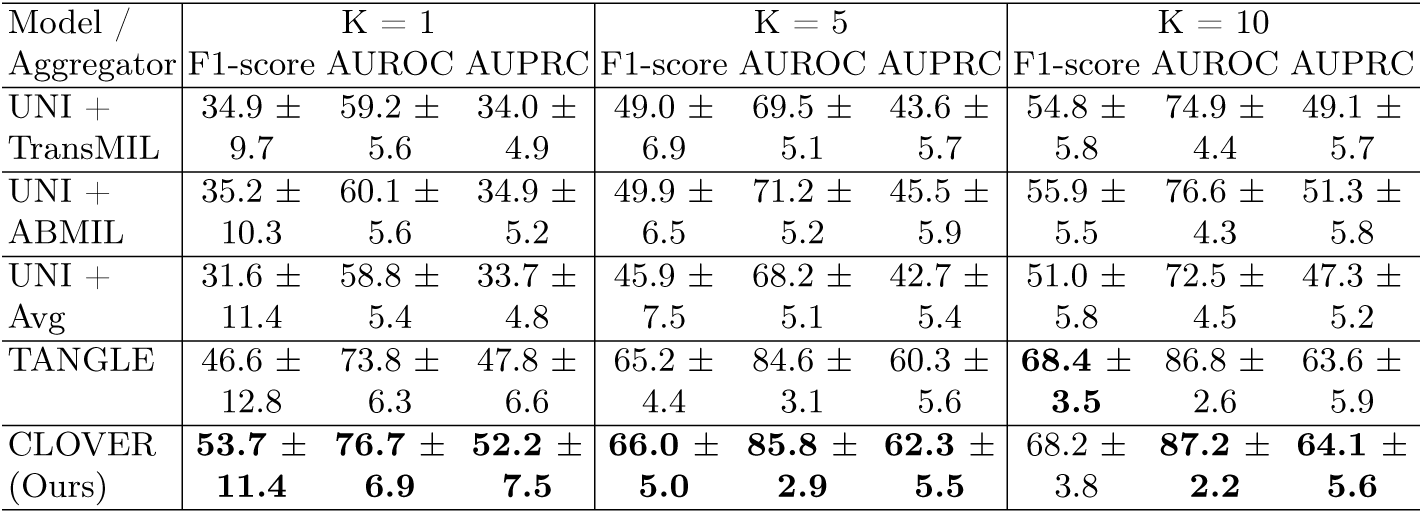
Prototype-based Molecular Subtype Classification in Human Breast Cancer. Performance comparison of different models and aggregation methods for classifying breast cancer molecular subtype (PAM50) classification across different few-shot settings (K=1,5,10), evaluated using F1-score, AUROC, and AUPRC metrics.

Despite the class imbalance in the PAM50 subtypes, CLOVER maintained robust performance across all metrics. At k=10, CLOVER achieved macro-AUPRC of 64.1 ± 5.6%, demonstrating effective discrimination between subtypes even with limited training data.

#### Clinical marker status prediction

The prediction of clinical markers (ER, PR, and HER2 status) represents a critical diagnostic task with direct therapeutic implications [60]. CLOVER demonstrated consistent improvements across all markers and k-shot scenarios (Table 3, 4, 5). For ER status prediction, performance scaled from 78.1 ± 18.3% (k=1) to 86.7 ± 5.5% (k=10) AUROC, consistently outperforming TANGLE and baseline methods. PR status prediction showed similar trends, with CLOVER achieving 68.4 ± 19.3% at k=1 and improving to 76.9 ± 6.5% at k=10.

**Table 3.** ER status prediction in human breast cancer. Performance comparison of different models and aggregation methods for predicting ER status across different few-shot settings (K=1,5,10), evaluated using F1-score, AUROC, and AUPRC metrics.

| Model /<br>Aggregator | K = 1 |  |  | K = 5 |  |  | K = 10 |  |  |
| --- | --- | --- | --- | --- | --- | --- | --- | --- | --- |
|  | F1-score | AUROC | AUPRC | F1-score | AUROC | AUPRC | F1-score | AUROC | AUPRC |
| UNI + | 58.7 ± | 59.4 ± | 81.2 ± | 68.3 ± | 71.1 ± | 87.1 ± | 71.1 ± | 75.5 ± | 89.2 ± |
| TransMIL | 12.6 | 10.6 | 6.9 | 8.1 | 8.7 | 5.1 | 6.6 | 7.2 | 4.1 |
| UNI + | 56.9 ± | 60.5 ± | 81.8 ± | 69.4 ± | 73.1 ± | 88.0 ± | 73.3 ± | 77.6 ± | 90.2 ± |
| ABMIL | 15.1 | 11.1 | 7.1 | 7.6 | 8.6 | 5.0 | 6.6 | 7.0 | 4.0 |
| UNI + | 54.0 ± | 59.5 ± | 81.1 ± | 66.8 ± | 70.7 ± | 86.9 ± | 70.4 ± | 75.1 ± | 89.1 ± |
| Avg | 16.9 | 10.5 | 6.9 | 8.2 | 8.6 | 5.0 | 6.8 | 7.3 | 4.0 |
| TANGLE | 63.4 ± | 75.3 ± | 88.4 ± | 79.2 ± | 85.0 ± | 93.0 ± | 80.9 ± | 86.3 ± | **93.6 ±** |
|  | 20.2 | 16.0 | 8.8 | 6.6 | 6.2 | 3.7 | 5.5 | 5.8 | 3.6 |
| CLOVER | **69.3 ±** | **78.1 ±** | **89.9 ±** | **80.2 ±** | **86.2 ±** | **93.3 ±** | **81.5 ±** | **86.7 ±** | 93.6 ± |
| (Ours) | 17.3 | 18.3 | 9.5 | 7.0 | 6.2 | 3.9 | 5.8 | 5.5 | 3.6 |

**Table 4.** PR status prediction in human breast cancer. Performance comparison of different models and aggregation methods for predicting PR status across different few-shot settings (K=1,5,10), evaluated using F1-score, AUROC, and AUPRC metrics.

| Model /<br>Aggregator | K = 1 |  |  | K = 5 |  |  | K = 10 |  |  |
| --- | --- | --- | --- | --- | --- | --- | --- | --- | --- |
|  | F1-score | AUROC | AUPRC | F1-score | AUROC | AUPRC | F1-score | AUROC | AUPRC |
| UNI + | 54.0 ± | 55.8 ± | 71.5 ± | 61.2 ± | 63.4 ± | 76.1 ± | 64.2 ± | 67.9 ± | 78.9 ± |
| TransMIL | 9.8 | 10.4 | 7.8 | 7.7 | 9.0 | 6.8 | 7.0 | 7.6 | 5.8 |
| UNI + | 53.0 ± | 56.6 ± | 72.0 ± | 62.0 ± | 65.0 ± | 77.1 ± | 65.7 ± | 69.6 ± | 79.9 ± |
| ABMIL | 11.6 | 10.9 | 7.9 | 7.7 | 8.9 | 6.5 | 7.4 | 7.6 | 5.5 |
| UNI + | 51.1 ± | 55.9 ± | 71.5 ± | 59.8 ± | 63.2 ± | 76.1 ± | 63.5 ± | 67.2 ± | 78.3 ± |
| Avg | 12.4 | 10.5 | 7.8 | 7.7 | 8.8 | 6.7 | 6.8 | 7.6 | 5.7 |
| TANGLE | 58.2 ± | 66.3 ± | 77.1 ± | 70.5 ± | 75.0 ± | 82.5 ± | 72.0 ± | 76.5 ± | **83.3 ±** |
|  | 16.6 | 17.9 | 11.2 | 8.0 | 8.3 | 5.8 | 6.5 | 6.6 | 5.1 |
| CLOVER | **64.2 ±** | **68.4 ±** | **78.3 ±** | **70.9 ±** | **75.9 ±** | **82.6 ±** | **72.6 ±** | **76.9 ±** | 83.0 ± |
| (Ours) | 15.9 | 19.3 | 11.5 | 8.3 | 7.6 | 5.5 | 7.0 | 6.5 | 5.1 |

**Table 5.** HER2 status prediction in human breast cancer. Performance comparison of different models and aggregation methods for predicting HER2 status across different few-shot settings (K=1,5,10), evaluated using F1-score, AUROC, and AUPRC metrics.

| Model /<br>Aggregator | K = 1 |  |  | K = 5 |  |  | K = 10 |  |  |
| --- | --- | --- | --- | --- | --- | --- | --- | --- | --- |
|  | F1-score | AUROC | AUPRC | F1-score | AUROC | AUPRC | F1-score | AUROC | AUPRC |
| UNI + | 51.8 ± | 51.2 ± | 22.4 ± | 54.0 ± | 53.3 ± | 23.9 ± | 56.4 ± | 55.5 ± | 25.4 ± |
| TransMIL | 12.7 | 10.2 | 7.1 | 9.3 | 10.2 | 8.0 | 7.1 | 9.7 | 8.7 |
| UNI + | 51.2 ± | 51.2 ± | 22.4 ± | 55.1 ± | 53.8 ± | 24.1 ± | 57.1 ± | 56.5 ± | 25.9 ± |
| ABMIL | 15.4 | 10.4 | 7.2 | 9.5 | 10.2 | 8.2 | 7.9 | 9.8 | 9.1 |
| UNI + | 50.1 ± | 51.0 ± | 22.1 ± | 55.6 ± | 53.0 ± | 23.5 ± | 57.6 ± | 55.7 ± | 25.1 ± |
| Avg | 18.2 | 10.4 | 7.1 | 9.7 | 10.0 | 7.8 | 7.4 | 9.4 | 7.9 |
| TANGLE | 49.4 ± | **54.9 ±** | **24.0 ±** | 58.7 ± | 58.9 ± | 27.4 ± | 61.3 ± | 61.5 ± | 30.3 ± |
|  | 19.9 | 10.4 | 8.6 | 10.5 | 9.4 | 9.2 | 8.4 | 8.6 | 9.9 |
| CLOVER | **51.1 ±** | 54.4 ± | 23.6 ± | **60.4 ±** | **60.8 ±** | **28.6 ±** | **63.8 ±** | **63.9 ±** | **31.1 ±** |
| (Ours) | 13.9 | 11.1 | 9.4 | 9.2 | 10.5 | 11.3 | 8.0 | 10.2 | 11.6 |

Notably, while HER2 status prediction proved more challenging across all methods, CLOVER still maintained a performance advantage, improving from 54.4 ± 11.1% at k=1 to 63.9 ± 10.2% at k=10. CLOVER consistently achieved higher F1-scores, particularly in low-shot scenarios, demonstrating robust generalization capability even with minimal labeled data.

### 4.4 Comparative analysis of multi-omics integration in CLOVER

To systematically evaluate the effectiveness of different omics combinations in CLOVER, we conducted comprehensive ablation studies across multiple classification tasks (Supplementary Figure 2). Our analysis revealed a clear hierarchy in the contribution of different omics modalities and their combinations to model performance. Among single-omics models, CNV demonstrated the lowest performance across all tasks with an average rank of 6.93, indicating its limited capability in isolation. MET and RNA showed relatively stronger individual predictive power, achieving average ranks of 2.87 and 4.73 respectively (Supplementary Figure 2A). This disparity suggests that methylation and transcriptomics data carry more directly relevant information for cancer classification tasks.

The integration of multiple omics modalities markedly improved model performance, with dual combinations showing substantial gains over single omics approaches. The RNA+MET combination proved particularly effective, achieving an average rank of 2.67 and demonstrating strong performance in both histologic subtype classification (k=10, AUROC: 90.0 ± 4.7) and PAM50 molecular subtype prediction (k=10, AUROC: 87.2 ± 1.8) (Supplementary Figure 2B, 2C). This synergy suggests that transcriptomic and methylation data capture complementary aspects of cancer biology.

Most notably, the full integration of all three modalities (RNA+MET+CNV) achieved the highest performance across tasks with an average rank of 2.60 (Supplementary Figure 2A). As shown in Figure 2B and 2C, this comprehensive integration maintained superior performance across different k-shot settings in both histologic and molecular subtype classification tasks. The advantage of full integration was particularly evident in challenging scenarios such as clinical marker prediction (Supplementary Figure 2D), where the model showed robust performance even in HER2 status determination (k=10, AUROC: 63.9 ± 10.2). These results demonstrate that while individual omics data provide valuable but partial perspectives on cancer biology, their integration enables the capture of complementary molecular features crucial for accurate classification. Notably, even CNV data, despite its lower individual performance, contributes meaningfully to the model’s predictive power when combined with other modalities. This suggests that CLOVER’s architecture effectively leverages the strengths of each omics type while mitigating their individual limitations through strategic integration.

### 4.5 Interpretability

To understand how CLOVER integrates information across modalities, we performed comprehensive interpretability analyses at both histological and molecular levels (Figure 3A). Our analysis revealed that CLOVER’s attention mechanisms effectively identify biologically significant regions while capturing meaningful molecular features associated with tumor characteristics.

**Fig. 3.**
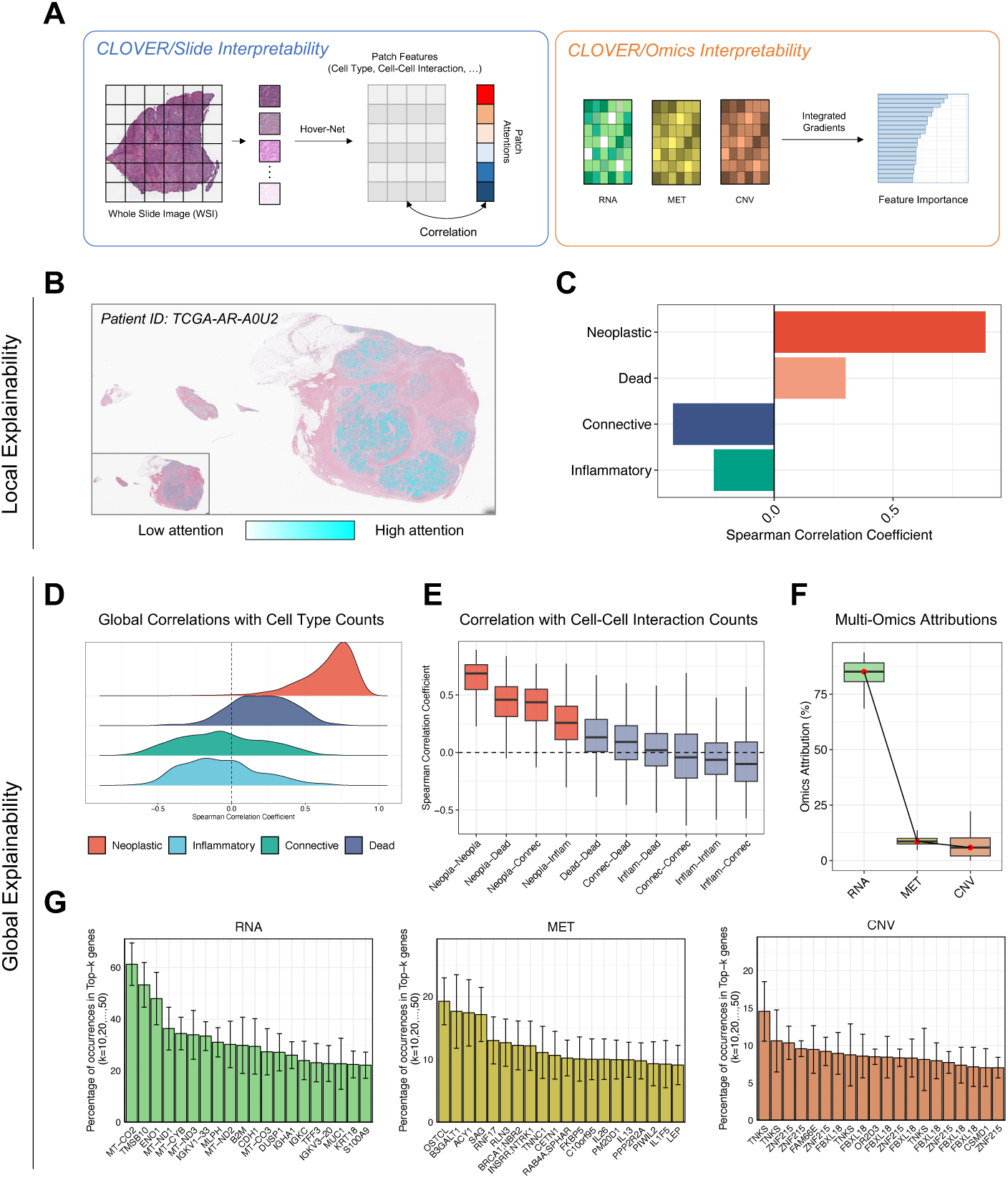
Multi-level interpretability analysis of CLOVER. (A) Overview of CLOVER’s dual interpretability framework, showing slide-level analysis using attention scores and patch features (left) and omics-level analysis using integrated gradients (right). (B) Representative whole slide image visualization showing attention heatmap overlay, with high attention regions highlighted in cyan and low attention regions in white. (C) Local correlation analysis between attention scores and cell type proportions in a single patient sample, demonstrating strong positive correlation with neoplastic cells and negative correlation with normal tissue components. (D) Global correlation analysis across all samples showing the distribution of Spearman correlation coefficients between attention scores and cell type counts per patch. (E) Global correlation analysis of cell-cell interaction counts per patch. (F) Attribution analysis showing relative contributions of different omics modalities to CLOVER’s predictions. (G) Feature attributions for each omics modality showing the most frequently selected features across samples, with error bars representing variation across different top-k thresholds (k=10,20,…,50).

At the slide level, we analyzed CLOVER’s patch-wise attention scores and their relationship with cellular composition using the Hover-Net cell detection framework. Examination of attention heatmaps from representative cases demonstrated that CLOVER naturally learns to focus on tumor-rich regions without explicit supervision (Figure 3B). Correlation analysis showed a strong positive correlation (*ρ* = 0.89) between attention scores and neoplastic cell density, while showing negative correlations with normal tissue components like connective (*ρ* = -0.43) and inflammatory cells (*ρ* = -0.25) (Figure 3C).

This pattern was consistent across the entire cohort, with global analysis revealing significantly higher attention correlation for neoplastic cells (median *ρ* = 0.70) compared to other cell types (Figure 3D). Notably, our analysis of tumor microenvironment features showed that CLOVER captures not only regions of high tumor cell density (median *ρ* = 0.69 for neoplastic-neoplastic interactions) but also areas of tumor-stroma interaction, with elevated attention scores in regions showing neoplastic-connective (*ρ* = 0.44) and neoplastic-inflammatory cell interactions (*ρ* = 0.26) (Figure 3E).

At the molecular level, integrated gradients analysis revealed that transcriptomic features contributed most significantly to the modality alignment (85.1% median attribution), followed by DNA methylation (8.6%) and copy number variations (5.9%) (Figure 3F). Detailed examination of feature importance across omics layers revealed biologically significant patterns that align with known cancer mechanisms (Figure 3G). Attributions of transcriptomic features highlighted several key functional groups of genes with known roles in breast cancer biology. Mitochondrial respiratory genes (MT-CO2, MT-ND1, MT-CYB, MT-ND3, MT-ND2, MT-CO3) emerged as highly important features, consistent with the suppression of mitochondrial gene expression in cancer cells and subsequent metabolic reprogramming [61]. The model also identified immune-related genes, particularly immunoglobulin genes (IGKV1-33, IGHA1, IGKC) associated with tumor-infiltrating lymphocytes and improved prognosis [62–65]. Additionally, CLOVER recognized tumor progression factors (TMSB10, ENO1), which are upregulated in breast cancer and promote tumor cell proliferation, migration, and invasion [66,67].

DNA methylation analysis highlighted several key epigenetic patterns, particularly in BRCA1-associated regions. BRCA1 promoter methylation, known to influence chemotherapy sensitivity, emerged as an important feature [68–71]. Copy number variation analysis, while showing lower overall attribution, identified alterations in regions containing TNKS, which influences Wnt signaling pathway regulation in breast cancers [72,73], and CSMD1, a putative tumor suppressor gene [74].

This multi-omic characterization demonstrates CLOVER’s ability to identify and integrate key molecular features that align with established cancer biology. The model’s spontaneous focus on these features suggests it has captured meaningful biological relationships between tissue morphology and molecular characteristics.

## 5 Discussion

In this study, we introduced CLOVER, demonstrating that multi-omics guided representation learning can significantly advance computational pathology analysis. Our comprehensive evaluation revealed that CLOVER successfully generates high-quality slide representations that effectively capture both morphological and molecular characteristics of tumors.

By comparing cohesion across histologic subtypes, molecular subtypes, and TNBC subtype, we demonstrated that our multi-omics guided approach learns more biologically meaningful representations compared to single-omics or image-based methods. Furthermore, our few-shot learning experiments validate CLOVER’s practical utility in limited-data scenarios, which is crucial for clinical applications, particularly significant for rare cancer subtypes where large labeled datasets are difficult to obtain. The model’s robust performance across different classification tasks with as few as 1-10 samples per class suggests that the learned representations effectively capture generalizable features of tumor biology.

Also, CLOVER’s ability to predict molecular characteristics directly from H&E slides could have significant clinical impact. The model’s performance in predicting ER, PR, and HER2 status suggests potential applications in rapid preliminary screening before traditional molecular testing, and resource-limited settings where molecular testing may be less accessible.

Our interpretability analysis further validates the biological relevance of CLOVER’s approach, revealing strong correlations between attention scores and tumor-rich regions, coupled with the identification of known cancer-associated genes and pathways through integrated gradients analysis.

Several important limitations must be acknowledged. Our evaluation relied solely on TCGA-BRCA data, which may not fully account for image quality variations across different institutions and imaging devices. Also, the lack of external validation across different institutions and patient populations limits our understanding of the model’s generalizability. While multi-omics integration improves performance, the current architecture may not optimally capture the complex hierarchical relationships between different molecular layers. Additionally, establishing causal relationships between morphological and molecular features remains challenging with current interpretability methods.

These limitations suggest promising future directions. Extending CLOVER to pan-cancer analysis could reveal shared biological mechanisms, though this requires careful consideration of tissue-specific variations. Future work should explore more sophisticated integration methods that can capture hierarchical relationships between molecular features, account for varying reliability and completeness of different omics data types, and incorporate prior biological knowledge about molecular interactions. Recent advances in multi-omics integration tools, such as graph neural networks, could enhance the capture of complex molecular interactions [75,76]. Integration with emerging spatial omics data could provide more direct validation of morphological-molecular relationships [77]. Future validation across multiple institutions is needed to assess performance across diverse imaging conditions and patient populations.

CLOVER represents a significant advancement in computational pathology by effectively bridging morphological and molecular features while maintaining interpretability. By addressing these limitations, future research can further advance toward more accurate and personalized cancer diagnosis and treatment.

