## Supplementary figures and images for "Contrastive Learning for Omics-guided Whole-slide Visual Embedding Representation"

### Supplementary Figure 1

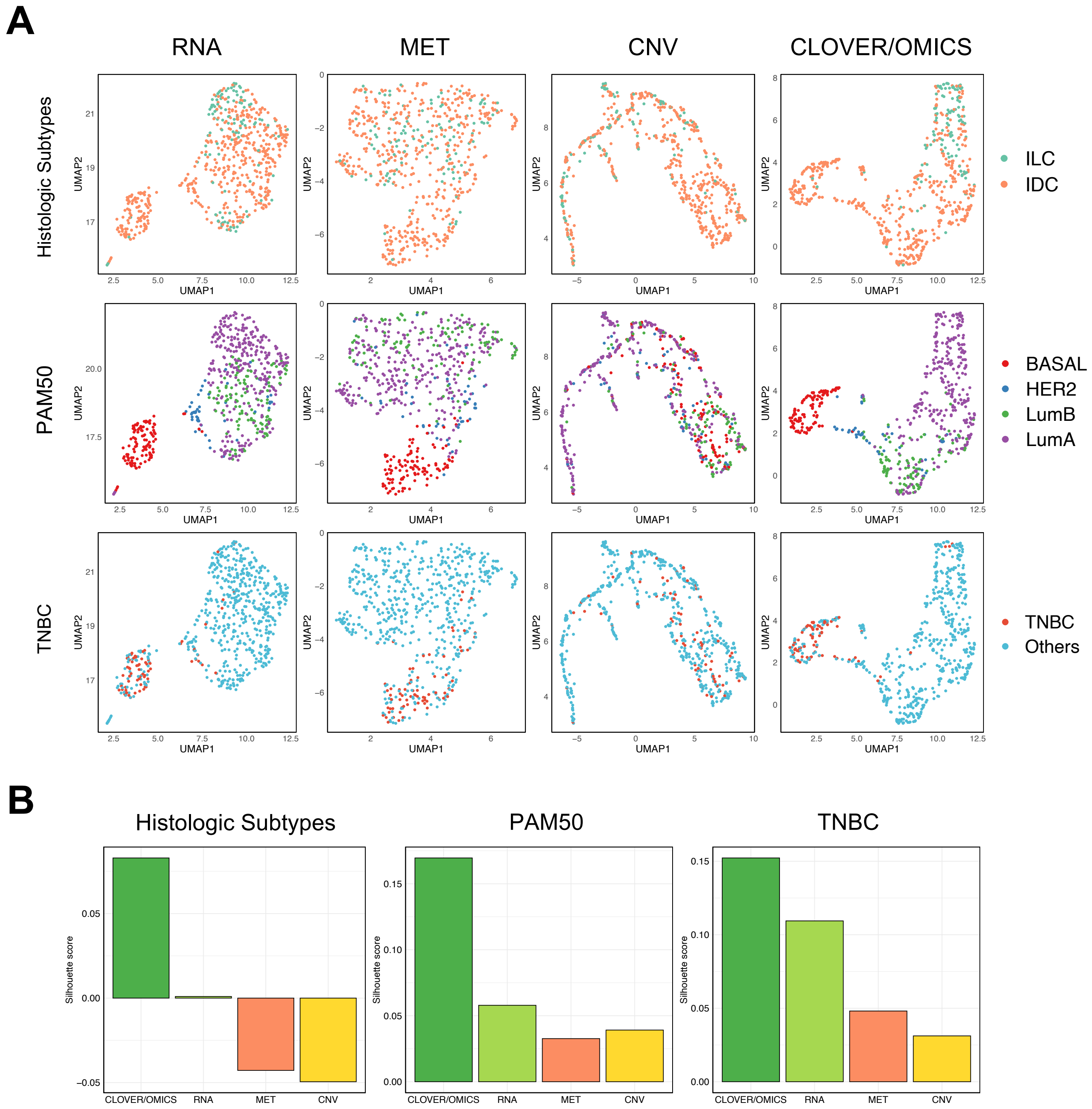
