## Supplementary Figure 2 for "Contrastive Learning for Omics-guided Whole-slide Visual Embedding Representation"

**A**

| | Avg Rank |
|---|---|
| RNA + MET + CNV | 2.60 |
| RNA + MET | 2.67 |
| RNA + CNV | 3.80 |
| MET + CNV | 3.87 |
| RNA | 4.73 |
| MET | 2.87 |
| CNV | 6.93 |

Average Rank: 1 2 3 4 5 6 7

**B**

Histologic Subtype

| | K=1 | K=5 | K=10 |
|---|---|---|---|
| RNA + MET + CNV | 80.7 ± 16.1 (1) | 88.1 ± 5.6 (2) | 89.9 ± 4.8 (3) |
| RNA + MET | 80.6 ± 16.3 (2) | 88.2 ± 5.7 (1) | 90.0 ± 4.7 (2) |
| RNA + CNV | 79.8 ± 15.7 (4) | 87.3 ± 5.5 (5) | 89.3 ± 4.8 (5) |
| MET + CNV | 75.3 ± 16.0 (6) | 85.9 ± 5.9 (6) | 88.5 ± 4.8 (6) |
| RNA | 80.0 ± 15.7 (3) | 87.6 ± 5.7 (4) | 89.4 ± 4.8 (4) |
| MET | 78.6 ± 15.4 (5) | 87.8 ± 5.6 (3) | 90.2 ± 4.3 (1) |
| CNV | 71.7 ± 18.1 (7) | 81.8 ± 6.6 (7) | 84.5 ± 5.4 (7) |

Rank: 1 2 3 4 5 6 7

**C**

Molecular Subtype (PAM50)

| | K=1 | K=5 | K=10 |
|---|---|---|---|
| RNA + MET + CNV | 76.7 ± 6.9 (2) | 85.8 ± 2.9 (1) | 87.2 ± 2.2 (1) |
| RNA + MET | 76.5 ± 7.0 (3) | 85.8 ± 2.7 (1) | 87.2 ± 1.8 (1) |
| RNA + CNV | 76.3 ± 6.9 (4) | 85.6 ± 3.0 (3) | 87.0 ± 2.3 (3) |
| MET + CNV | 76.3 ± 7.7 (4) | 84.6 ± 3.2 (5) | 86.1 ± 1.9 (6) |
| RNA | 74.7 ± 6.6 (6) | 84.5 ± 2.9 (6) | 86.2 ± 2.0 (4) |
| MET | 77.2 ± 6.6 (1) | 84.9 ± 3.1 (4) | 86.2 ± 2.5 (4) |
| CNV | 67.0 ± 8.8 (7) | 77.8 ± 4.9 (7) | 81.0 ± 3.6 (7) |

Rank: 1 2 3 4 5 6 7

**D**

Clinical Markers

| | ER K=1 | ER K=5 | ER K=10 | PR K=1 | PR K=5 | PR K=10 | HER2 K=1 | HER2 K=5 | HER2 K=10 |
|---|---|---|---|---|---|---|---|---|---|
| RNA + MET + CNV | 78.1 ± 18.3 (3) | 86.2 ± 6.2 (4) | 86.7 ± 5.5 (4) | 68.4 ± 19.3 (4) | 75.9 ± 7.6 (4) | 76.9 ± 6.5 (4) | 54.4 ± 11.1 (3) | 60.8 ± 10.5 (2) | 63.9 ± 10.2 (1) |
| RNA + MET | 78.1 ± 18.2 (3) | 86.5 ± 6.2 (3) | 86.9 ± 5.6 (3) | 68.8 ± 19.5 (3) | 76.0 ± 8.0 (3) | 77.0 ± 6.7 (3) | 53.6 ± 9.9 (5) | 59.3 ± 9.8 (4) | 63.3 ± 9.4 (3) |
| RNA + CNV | 77.7 ± 18.5 (5) | 86.1 ± 6.2 (5) | 86.7 ± 5.5 (4) | 68.4 ± 18.9 (4) | 75.8 ± 8.0 (5) | 76.6 ± 6.4 (5) | 54.6 ± 11.4 (2) | 61.3 ± 10.5 (1) | 63.8 ± 9.6 (2) |
| MET + CNV | 79.0 ± 19.8 (2) | 86.9 ± 6.2 (1) | 87.3 ± 5.4 (1) | 69.1 ± 19.8 (1) | 77.0 ± 7.2 (1) | 77.8 ± 5.8 (1) | 52.1 ± 9.8 (7) | 58.0 ± 9.6 (6) | 61.9 ± 9.3 (5) |
| RNA | 75.7 ± 17.5 (6) | 85.0 ± 6.7 (6) | 85.7 ± 6.0 (6) | 67.3 ± 18.1 (6) | 74.7 ± 8.6 (6) | 76.0 ± 7.0 (6) | 54.7 ± 10.9 (1) | 60.3 ± 9.9 (3) | 63.1 ± 9.4 (4) |
| MET | 80.0 ± 18.7 (1) | 86.7 ± 6.2 (2) | 87.2 ± 5.7 (2) | 69.1 ± 19.9 (1) | 76.7 ± 7.4 (2) | 77.5 ± 6.0 (2) | 53.9 ± 10.2 (4) | 58.6 ± 10.2 (5) | 61.5 ± 9.5 (6) |
| CNV | 64.8 ± 18.1 (7) | 77.9 ± 8.6 (7) | 81.0 ± 6.4 (7) | 60.0 ± 16.1 (7) | 68.6 ± 9.0 (7) | 72.1 ± 6.6 (7) | 52.2 ± 10.7 (6) | 55.9 ± 10.1 (7) | 59.2 ± 10.1 (7) |

Rank: 1 2 3 4 5 6 7
